# Immunity gene silencing increases transient protein expression in *Nicotiana benthamiana*

**DOI:** 10.1101/2025.01.30.635616

**Authors:** Isobel L. Dodds, Emma C. Watts, Mariana Schuster, Pierre Buscaill, Yasin Tumtas, Nicholas J. Holton, Shijian Song, Johannes Stuttmann, Matthieu H. A. J. Joosten, Tolga O. Bozkurt, Renier A. L. van der Hoorn

**Affiliations:** The Plant Chemetics Laboratory, Department of Biology, University of Oxford, South Park Road, OX1 3RB Oxford, UK; Department of Life Sciences, Imperial College London, London, UK; Leaf Expression Systems, Norwich Research Park, Colney Lane, NR4 7UJ, Norwich, UK; Aix Marseille University, CEA, CNRS, BIAM, UMR7265, LEMiRE (Microbial Ecology of the Rhizosphere), 13115, Saint‑Paul lez Durance, France; Laboratory of Phytopathology, Wageningen University and Research, Droevendaalsesteeg 1, 6708 PD Wageningen, The Netherlands

## Abstract

Agroinfiltration of *Nicotiana benthamiana* is routinely used for transient gene expression in plant sciences and molecular pharming. Here, we depleted transcripts of 21 different immunity-related genes through virus-induced gene silencing (VIGS) to identify genes that hamper transient expression in juvenile plants. These experiments uncovered that silencing of *ethylene insensitive- 2* (*EIN2*), receptor-like kinase *CERK1*, transcription factor regulator *nonexpressor of pathogenesis-related genes 1* (*NPR1*) and *isochorismate synthase* (*ICS*), increases transient GFP expression by 2-, 4-, 4- and 11-fold, respectively. Accordingly, the *npr1a/npr1b* double mutant of *N. benthamiana* does indeed facilitate increased protein accumulation when transiently expressed. These results indicate that glycan perception through CERK1, and ET and SA signaling pathways via EIN2 and NPR1/ICS, respectively, contribute to suppressed transient gene expression in frequently used juvenile *N. benthamiana* plants.

---

The infiltration of *Nicotiana benthamiana* with *Agrobacterium tumefaciens* (agroinfiltration) has become a routine expression platform for plant science and molecular pharming, yet this platform remains to be further optimized. We recently showed that *N. benthamiana* silenced for the *cold shock protein* (*CSP*) *receptor* (*CORE*) enables 8-fold more GFP production in older, 6–8-week-old plants, which are normally not used because of low transient expression efficiencies (Dodds et al., 2023). Here, we investigated whether we can also increase transient protein expression levels in routinely used younger, 5-week-old juvenile plants, by silencing immunity-related genes.

We selected 21 immunity-related genes encoding proteins that act at different levels in the plant immune system (Supplemental **Table S1**). Besides *CORE*, we silenced receptor-encoding genes *WAK1* (*Wall-associated Protein Kinase*), *CERK1* (*Chitin Elicitor Receptor Kinase-1*), *BAK1* (*BRI1-associated Receptor Kinase-1*), *SOBIR1* (*Suppressor of BIR1-1*) and *RE02* (*Receptor of SCPs*). We also tested silencing of immune signaling components such as *F-box protein ACIF1* (*Avr/Cf-induced F-box-1*); lipase-like proteins *EDS1* (*Enhanced Disease Susceptibility-1*) and *SAG101* (*Senescence-associated Gene-101*); *CDPK* (*Calcium-dependent Protein Kinase*), *MPK3/6* (*MAP protein kinases-3 and -6*), *Nod-like helper receptors NRC2/3/4* (*NLR Required for Cf Signaling*) and chaperone *CRT3a* (*Calreticulin-3a*). We also included genes required for stress hormone signaling, including *PAL* (*Phenylalanine Ammonia Lyase*), *ICS* (*Isochorismate Synthase*), *NPR1* (*Nonexpressor of PR genes-1*), *EIN2* (*Ethylene-insensitive-2*) and *WRKY* transcription factors. Finally, we included genes encoding *AHA2* (*Arabidopsis H*^*+*^*-ATPase 2*) and *RBOHB* (*Respiratory Burst Oxidase Homolog B*). Genes encoding phytoene desaturase (*PDS*) and *ß-glucuronidase* (*GUS*) were included as positive and negative controls for silencing, respectively. We resynthesized the silencing fragments as published previously (Supplemental **Table S1**) and selected novel fragments targeting *ACIF1, CDPK, CORE, ICS* and *AHA2* (Dodds et al. 2023, Supplemental **Tables S2** and **S3**).

Tobacco Rattle Virus (TRV) vectors, each carrying a fragment of these 21 immunity genes and the controls were agroinfiltrated into 2-week-old seedlings and plants were tested for transient expression three weeks later. Transcript levels of the targeted genes were downregulated with novel silencing fragments (Supplemental **Figure S1**). At that stage, no strong phenotypes were observed in TRV-inoculated plants, except for photobleaching in *TRV::PDS* plants, dwarfed *TRV::BAK1* and *TRV::CDPK* plants, and small, chlorotic *TRV::ICS* plants (Supplemental **Figure S2**). Leaf discs of *TRV::RBOHB* and *TRV::BAK1* plants showed a reduced oxidative burst upon flg22 treatment (Supplemental **Figure S3**), consistent with effective silencing of these genes.

Silenced plants were agroinfiltrated with a 1:1:1 mixture of Agrobacteria delivering three ‘traffic light’ reporters: bioluminescent AgroLux bacteria (Jutras et al., 2021, orange), mixed with Agrobacteria delivering expression cassettes for cytonuclear GFP (cGFP, green) and secreted RFP (sRFP, red). GFP fluorescence in leaf discs taken at 5 days-post-infiltration (5dpi) was significantly higher in *TRV::ICS, TRV::CERK1, TRV::NPR1* and *TRV::EIN2* plants, when compared to *TRV::GUS* plants (**Figure 1A**). The same silenced plants also showed more RFP fluorescence when compared to *TRV::GUS* control plants (Supplemental **Figure S4A**). No altered fluorescence was detected in *TRV::CORE* plants, consistent with low *CORE* expression in juvenile plants (Wang et al., 2016). AgroLux bioluminescence was similar between these silenced plants (Supplemental **Figure S4B**), indicating that silencing these immunity genes does not increase Agrobacterium population levels. Scanning of agroinfiltrated leaves in subsequent experiments confirmed that silencing *CERK1* or *NPR1* increased GFP-fluorescence by >4-fold and *ICS* silencing even >10-fold, when compared to the *GUS* silencing control (**Figure 1B and 1C**). Also a significant, 1.7-fold increased GFP fluorescence was detected in *TRV::EIN2*, when compared to *TRV::GUS* plants (Supplemental **Figure S5**), consistent with the initial screen.

**Figure 1.**
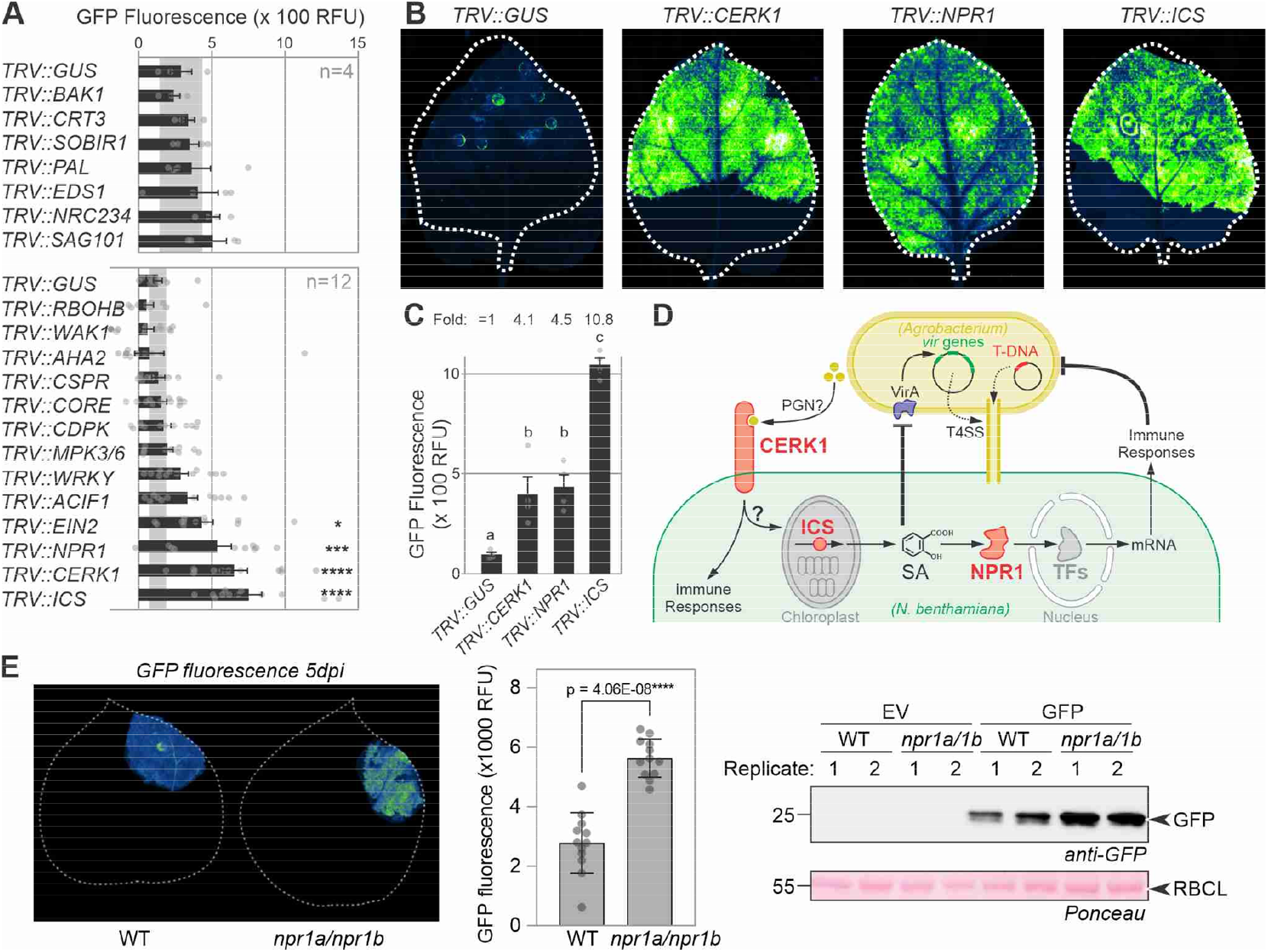
Immunity gene silencing increases transient expression in *Nicotiana benthamiana* **(A)** Two-week old *N. benthamiana* plants were inoculated with Tobacco Rattle Virus (TRV) carrying fragments of 21 immunity genes in two different experiments. Systemic leaves were agroinfiltrated 3 weeks later with a 1:1:1 mixture of Agrobacteria carrying binary vectors for expressing cytoplasmic GFP and secreted RFP, and bioluminescent AgroLux bacteria. Leaf discs were taken at 5 dpi and analyzed for GFP fluorescence. Error bars represent SE of n=4 (top) and n=12 (bottom) replicates. Data were analyzed by ANOVA with Dunnett’s post hoc test. *, p<0.05; **, p<0.01; ****, p<0.0001. **(B)** Silencing of *CERK1, NPR1* and *ICS* increases transient GFP expression. 2-week old *N. benthamiana* plants were inoculated with Tobacco Rattle Virus (TRV) carrying fragments of various genes to be targeted. Plants were agroinfiltrated 3 weeks later with Agrobacterium carrying a binary vector for GFP expression, and images were taken at 5dpi and quantified. **(C)** Quantification of GFP fluorescence detected in (B). Error bars represent SE of n=4 replicates. ANOVA with a Dunnett’s Multiple Comparison test. **(D)** Likely roles of CERK1, ICS and NPR1 in suppressing transient gene expression. CERK1 is a receptor-like kinase that might perceive peptidoglycan (PGN). Isochorismate synthase (ICS) is a metabolic enzyme producing a precursor for salicylic acid (SA) in the chloroplast. Non- expressor of PR1 (NPR1) regulates transcription factors (TFs) that activate immunity genes. **(E)** Increased transient GFP expression in the *npr1a/npr1b* double mutant. GFP was transiently expressed without P19 and fluorescence was imaged at 5 dpi (left), quantified (middle) and leaf extracts were analysed for anti-GFP western blot (right). Error bars represent SD of n=18 replicates.

The observed effects imply that immunity genes *CERK1, NPR1* and *ICS* encode important barriers for transient expression (**Figure 1D**). CERK1 is a receptor-like kinase involved in the perception of chitin (Mya et al., 2007) and peptidoglycan (Willmann et al., 2011). NPR1 is a transcriptional regulator of responses induced by salicylic acid (SA) (Spoel & Dong, 2024). ICS is a chloroplastic enzyme producing isochorismate, a precursor of SA (Spoel & Dong, 2024). The unaltered GFP fluorescence in *TRV::PAL* plants indicates that SA restricting transient expression is not produced via phenylalanine, but through isochorismate in agroinfiltrated leaves (Spoel & Dong, 2024). The effect of *ICS* silencing is twice as strong compared to *NPR1* silencing might be because ICS depletion would not only reduce immune signaling, but also reduce SA levels that negatively regulate *vir* gene expression in Agrobacterium (Yuan et al., 2007).

To enhance transient expression without relying on VIGS, we are disrupting the open reading frames of these immunity-related genes using genome editing and stacking mutant genes. So far, the deletion of the two silenced *NPR1* genes (Supplemental **Figure S6**), has indeed caused increased GFP fluorescence and accumulation (**Figure 1E**), and enhanced IgG accumulation (Supplemental **Figure S7**). These genome-edited lines hold important potential for boosting transient gene expression for protein production in *N. benthamiana*.

## Acknowledgements

We thank Urszula Pyzio for excellent plant care, and Sarah Rodgers, Caroline O’Brian and Patricia Bowman for technical support.

## Funding

This project was financially supported by the Interdisciplinary DTC project DDT00060 (ID) and DDT00230 (EW); BBSRC projects BB/R017913/1 (PB) and BB/S003193/1 (MS); and ERC project 101019324 (RH).

## Author contributions

RH conceived the project; ID and EW performed most experiments with help of MS and PB and SS; YT, MJ and TB provided VIGS constructs and JS provided the *npr11/npr1b* mutant; ID, EW and RH wrote the manuscript with all authors.

## Competing interests

none declared

## Notes

### Competing Interest Statement

The authors have declared no competing interest.

