## Supplemental for "Immunity gene silencing increases transient protein expression in *Nicotiana benthamiana*"

### Materials and Methods

**Plant growth -** *N. benthamiana* plants (LAB strain) were grown at a constant temperature of 21°C and approximately 60% relative humidity under a 16-hour light (40 µmol $m^{-2}$ $s^{-1}$) and 8-hour dark regime.

**Cloning silencing constructs** – All used plasmids are listed in Supplemental **Table S2**. Fragments for silencing *CORE, ACIF1, ICS, CDPK* and *AHA2* were designed using the SolGenomics VIGS tool (Fernandez-Pozo et al., 2015) and are listed in Supplemental **Table S3**. All other silencing fragments were selected from the literature and resynthesized and are listed in Supplemental **Table S3**. Silencing fragments were cloned into TRV2gg (Duggan et al., 2021) by Golden Gate cloning using the BsaI reaction, resulting in the binary plasmids listed in Supplemental **Table S2**. Plasmids were transformed into *E. coli* DH10β or DH5α and clones carrying inserts were selected by colony PCR. Plasmids were purified and the inserts verified by sequencing. Binary plasmids were transformed into Agrobacterium GV3101-pMP90 and transformants were selected on LB agar plates containing 25 mg/L rifampicin, 10 mg/L gentamycin and 50 mg/L kanamycin. A single transformant colony was cultured in liquid LB containing the same antibiotics.

**Cloning cGFP/sRFP reporters** – To create a binary plasmid that carries 35S:cGFP without the p19 silencing inhibitor, cGFP was amplified from pPJ048 (Jutras et al., 2021) using primers cGFPf and cGFPr (Supplemental **Table S2**) and cloned into pJK001c using a BsaI reaction, resulting in binary vector pID41 (35S:cGFP). To create a binary plasmid that carries 35S:sRFP without the p19 silencing inhibitor, sRFP was amplified from pPJ057 (Beritza et al., 2024) using primers sRFPf and sRFPr (Supplemental **Table S2**) and cloned into pJK001c using a BsaI reaction, resulting in binary vector pID42 (35S:sRFP).

**Virus-induced Gene Silencing (VIGS) -** Agrobacteria containing TRV1 or TRV2 (Supplemental **Table S2**) were grown overnight at 28°C in LB medium containing 25 mg/L rifampicin, 10 mg/L gentamycin and 50 mg/L kanamycin. The cultures were centrifuged at 3500 x *g* for 10 minutes at room temperature and the pellet was resuspended at $OD_{600}$ = 1.0 in agroinfiltration buffer (10 mM MES pH 5.7, 10 mM $MgCl_2$, 100 µM acetosyringone). TRV2 cultures were mixed at a 1:1 ratio with TRV1 cultures. Two-week-old *N. benthamiana* were agroinfiltrated with the bacterial suspension mixture using a 1 ml needleless syringe. Three to five weeks later plants were assessed for silencing by observing whether the *TRV::PDS* plants had bleached leaves. Silenced plants were then tested by agroinfiltration.

**Traffic Light Screen** – Plants inoculated with TRV silencing constructs were agroinfiltrated with a mixture of three cultures: Agrobacterium carrying an expression cassette for cGFP, Agrobacterium carrying an expression cassette for sRFP and Agrobacterium carrying luciferase (AgroLux, Jutras et al.,

2021). At 5 days post agroinfiltration, leaf discs (6 mm diameter) were punched and placed onto 200 µl sterile water in a white, opaque 96-well plate (Corning). Fluorescence and luminescence were detected using a plate reader. GFP fluorescence was detected with ex500nm/em530nm at gain 100 and RFP fluorescence was measured ex550nm/em585nm at gain 100. For each agroinfiltrated sector, at least two leaf discs (technical replicates) were taken to reduce the effect of variability in expression within leaves. Statistics were performed using a One-way ANOVA followed by Dunnett's multiple comparisons test using GraphPad Prism9 for macOS.

**Fluorescent protein leaf imaging and quantification -** Plants inoculated with silencing constructs for *GUS, CERK1, NPR1* and *ICS* were agroinfiltrated across 60-100% of a leaf with Agrobacterium carrying an expression cassette for cGFP. At 5dpa, leaves were detached and scanned on an Amersham Typhoon 5 Biomolecular Imager (GE Healthcare Life Sciences, Little Chalfont, UK), using the 488nm laser and Cy2 filter at PMT300 to detect *in planta* fluorescence at 5dpi. The ImageJ green fire blue LUT was applied to images. Quantification was performed using ImageJ. Data was analyzed using ANOVA with a Dunnett's Multiple Comparison test.

**RT-PCR** – 100 mg of leaf tissue was flash-frozen in liquid nitrogen and ground into a fine powder using a pestle and mortar. The Qiagen RNeasy Mini Kit (including the Qiagen Shredder columns) was used to extract total RNA. The extracted RNA was treated using Qiagen RNase-free DNase and purified using the Qiagen RNeasy Mini Kit. The RNA was measured using a NanoDrop. Reverse transcription (RT) was performed on 1µg RNA by following the first-strand cDNA synthesis using the SuperScript™ II RT protocol, using oligo-dT primers to produce cDNA. PCR was performed using Q5 polymerase with gene-specific primers. PCR products were separated on a 3% agarose gel and imaged using a G:Box imaging system. Quantification was performed using ImageJ. Data was statistically analyzed using an unpaired *t*-test using GraphPad Prism 9 for macOS.

**ROS assays -** ROS assays were performed as described previously (Buscaill et al., 2019). Briefly, after incubation in water overnight, *N. benthamiana* leaf discs (6 mm diameter) were transferred to a 96-well plate (one per well) and 100 µl of a solution containing 25 ng/µl luminol, 25 ng/µl Horse Radish Peroxidase (HRP) and 100 nM flg22, was added to each leaf disc. Chemiluminescence was measured immediately with the Infinite M200 plate reader (Tecan, Mannedorf, Switzerland) every minute for one hour. Standard errors were calculated at each time point and for each treatment.

**Generation of *npr1a/npr1b* mutant** – Putative homologs of Arabidopsis NPR1/3/4 in *N. benthamiana* were determined by reciprocal BLAST (tBLASTn) searches identifying first homologs from *Solanum lycopersicum*, and subsequently using the tomato proteins as queries to search within the *N. benthamiana* genome. Two genes encoding proteins most similar to AtNPR1 were designated *NbNPR1a* and *NbNPR1b*. sgRNA transcription units under Arabidopsis promoter control were generated in plasmids pDGE332, 333, 335, 337 as previously described (Stuttmann et al., 2021) for targeting the sequences *ATGAGGCAAGGCCTTATCAANGG*, *GAGCAGAACTTGGTCTACAANGG* and *AACATGTTAAGAGGATACATNGG* (PAM sequence underlined) present in both *NPR1a* and

*NPR1b*. The sgRNA transcription units were subsequently assembled in pDGE311, containing zCas9i under 35S promoter control (Grutzner et al., 2021), a kanamycin resistance cassette for positive selection and the CaBs3 gene for negative selection (Stuttmann et al., 2021). The resulting plasmid pDGE370 was electroporated into Agrobacterium strain GV3101 pMP90, and *N. benthamiana* was transformed as previously described (Ordon et al., 2019). Non-transgenic individuals from the T1 generation were genotyped using oligonucleotides JS1767/1768 (*NPR1a*; *TCCATTGCATATTTCCTTTGCAG* and *AGTGCAAGATCTAGAAGTTCTG*) and JS1769/1768 (*NPR1b*; *TTCCACTTCACATTTCCTTCCC* and *AGTGCAAGATCTAGAAGTTCTG*).

**Fluorescent protein leaf imaging and western blot -** Six-week-old WT and *npr1a/npr1b* plants were agroinfiltrated with OD600 = 0.5 Agrobacterium carrying pBM2 delivering sfGFP without p19 silencing inhibitor. At 5 dpi, leaves were scanned on the Amersham Typhoon using the Cy2 filter at PMT300. Four 6 mm leaf discs were taken from the same tissue and extracted into phosphate buffer (Merck PBS tablets) with 0.05% Tween-20 (PBS-T), 1 mM EDTA, and 5 µM Halt™ Protease Inhibitor Cocktail (ThermoFisher). Samples were centrifuged for 15 minutes at 15,000 *x* g, 4˚C after which the supernatant was extracted into a clean Eppendorf tube. Samples were mixed 1:1 with Laemmli buffer (0.125 M Tris, 0.14 M SDS, 20% glycerol, 10% 2-mercaptoethanol, 0.2% Bromophenol Blue) and denatured by boiling and loaded into 12% SDS-PAGE gels, running at 35 mA for 2 hours in Invitrogen Novex vertical gel tanks. Gels were then transferred onto a polyvinylidene difluoride (PVDF, BioRad) membrane using BioRad Trans-blot Turbo® (BioRad Kit 1704275). Blots were blocked for 1 hour at 21 °C with 5% (w/v) skim milk in PBS-T. The membrane was incubated with 1:5000 HRP-conjugated goat anti-GFP antibody (Abcam, Ab6663) in 5% (w/v) skim milk for one further hour at room temperature. Blots were washed twice with PBS-T for 5 minutes, and chemiluminescence was detected with Clarity Western ECL Substrate (BioRad) visualised using the ImageQuant® LAS-4000 imager (GE Healthcare, Healthcare Life Sciences, Little Chalfont, UK).

**ELISA of IgG antibody -** Six-week-old WT and *npr1#2* plants were agroinfiltrated with a 1:1:1:1 mix of Agrobacterium delivering silencing inhibitor p19 (pJK268c, Kourelis et al., 2020); sfGFP (pBM2); COVA2-15 LC (pKB21, Beritza et al., 2025) and COVA2-15 HC (pKB22, Beritza et al., 2025) (OD600 = 0.5) and total extract was collected as above. 15 ng of SARS-CoV-2 Spike Protein Ribosome Binding Domain (ThermoFisher RP-87678) in ELISA carbonate buffer ($Na_2CO_3$: 0.015 M, $NaHCO_3$: 0.035 M, $NaN_3$: 0.003 M) was added to wells of a 96-well ELISA plate (ThermoFisher 467120). The plate was sealed with plate sealer (ThermoFisher, 3501) and incubated overnight at 4˚C. Wells were washed 3 times with PBS-T, and 3 times with MilliQ before adding PBS-T with 3% skim milk (PBS-TM) and incubating for 2 hours at room temperature. Total extract of leaf samples was diluted into PBS-TM (1:3 – 1:3000) and added into the ELISA plate for a further 2-hour incubation. After repeating the wash step, HRP-conjugated mouse anti-human IgG Fc Antibody (GenScript A01854) was diluted to 1:20000 in PBS-TM and added to wells. The wash step was repeated once more before adding TMB substrate (ThermoFisher, 34028). After 5 minutes, the reaction was stopped with 1M HCl, and absorbance at 450nm was measured in a plate reader (Tecan Spark®).

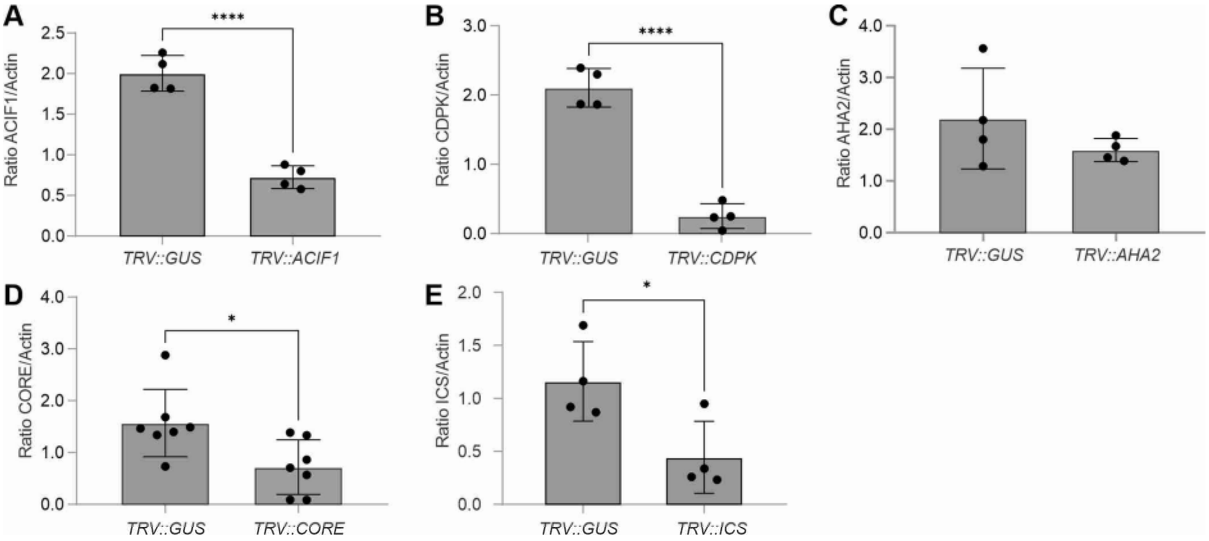


**Figure S1** Novel TRV constructs silence target genes.

RNA was extracted from systemic leaves of 5-week old plants, 3 weeks after introducing TRV constructs by agroinfiltration. cDNA was generated and used as a template for semi-quantitative PCR with gene-specific primers. Error bars represent SE of n=4 (A-C, E) or n=7 (D) replicates. Data were analyzed using unpaired *t*-tests (*, $p<0.05$; ****, $p<0.0001$).

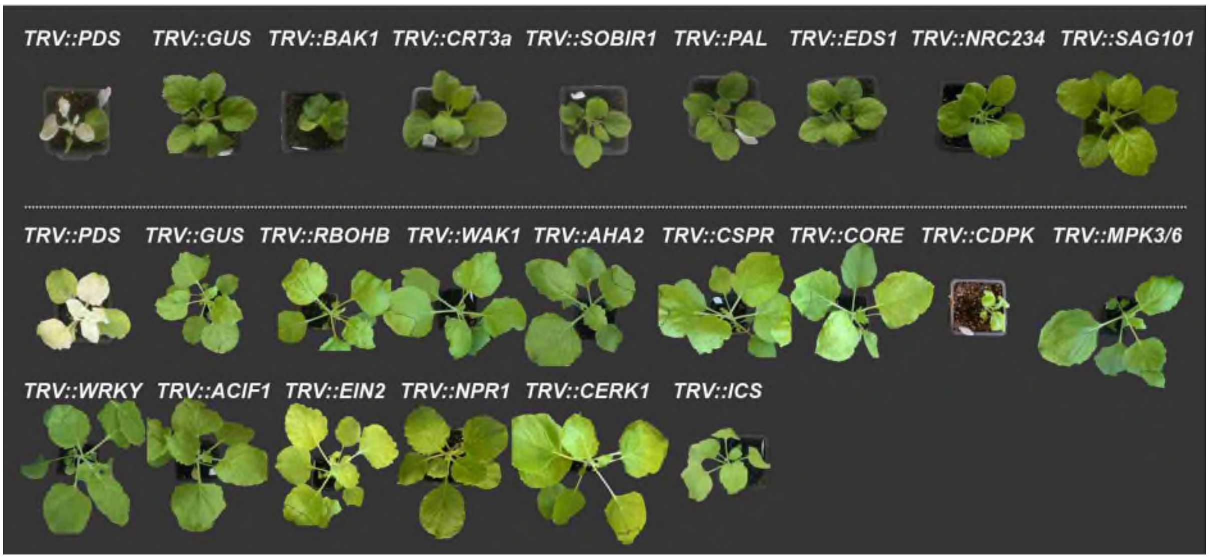


**Figure S2** Phenotypes of VIGSed plants

Pictures were taken when plants were 5 weeks old, 3 weeks after agroinoculating TRV. Plants and pots were cropped from the background and scaled to a similar pot size (6.5 x 6.5 cm).

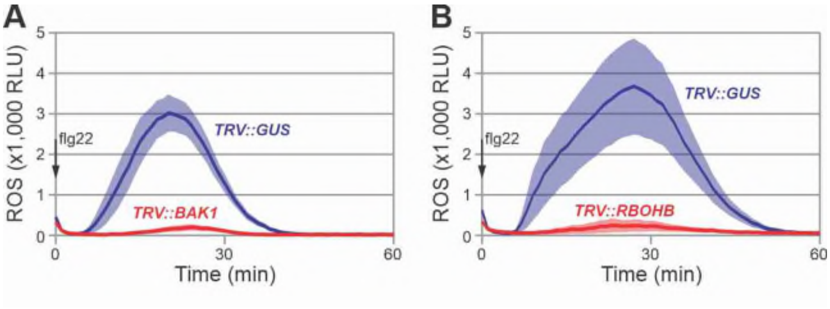


**Figure S3** Suppressed flg22-induced oxidative burst in *TRV::BAK1* and *TRV::RBOHB* plants. Leaf discs from TRV-inoculated plants floating on luminol HRP (Horse Radish Peroxidase) were treated with 100 nM flg22 and the release of reactive oxygen species (ROS) was measured for 60 minutes as bioluminescence in a plate reader. Error shades represent SE of n=6 replicates.

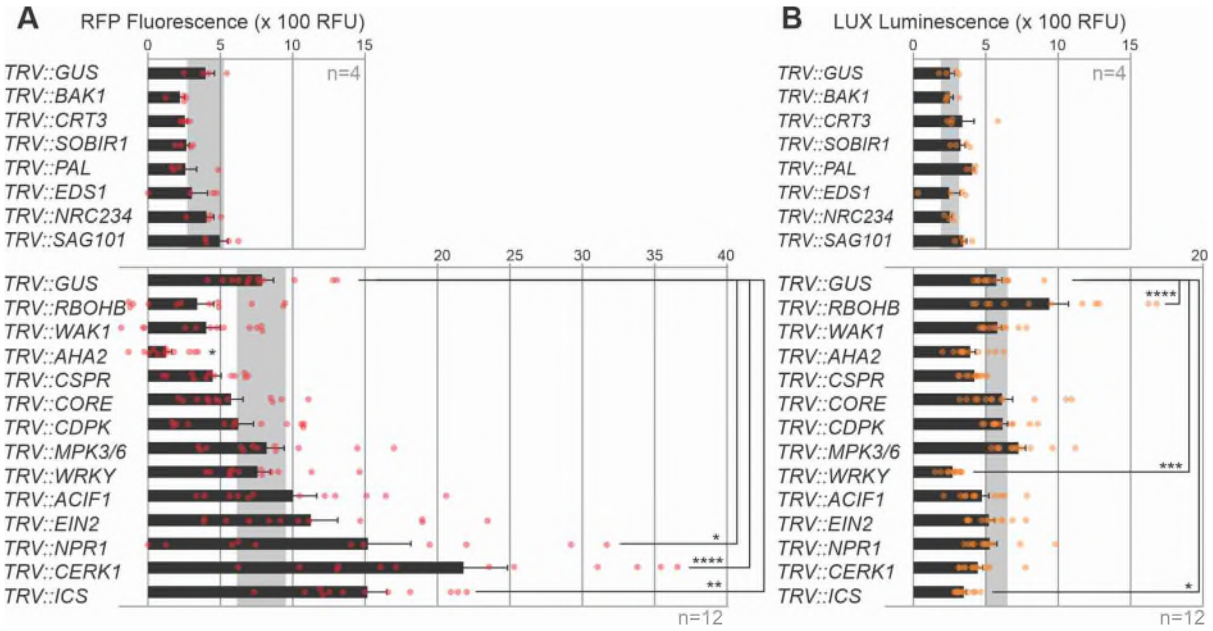


**Figure S4** Impact of immunity gene silencing on sRFP fluorescence and AgroLux luminescence.

2-week old *N. benthamiana* plants were agroinoculated with TRV carrying fragments of 21 immunity genes. Three weeks later, systemic leaves were agroinfiltrated with a 1:1:1 mixture of Agrobacteria carrying a binary vector for the expression of cGFP and sRFP, and bioluminescent AgroLUX bacteria. Leaf discs were taken at 5dpi and analyzed for fluorescence and bioluminescence. GFP fluorescence is shown in **Figure 1A**; sRFP fluorescence in **(A)** and AgroLux bioluminescence in **(B)**. The significant increase in bioluminescence in *TRV::RBOHB* plants was not detected in replicate experiments. Error bars represent SE of n=4 (top) and n=12 (bottom) replicates. Data were analyzed by ANOVA with Dunnett's post hoc test. *, $p<0.05$; **, $p<0.01$; ****, $p<0.0001$.

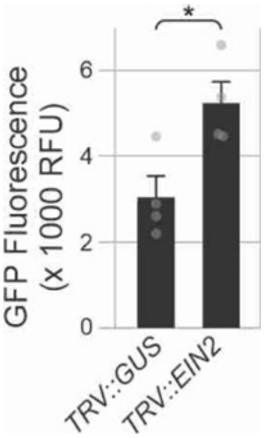


**Figure S5** EIN2 silencing increases transient expression levels.

2-week-old *N. benthamiana* plants were inoculated with *TRV::GUS* and *TRV::EIN2* and three weeks later, systemic leaves were agroinfiltrated for transient cGFP expression. Five days later, leaves were detached and scanned for GFP fluorescence. Error bars represent SE of n=4 replicates. Significance was determined with a Students *t*-test (*, p=0.020).

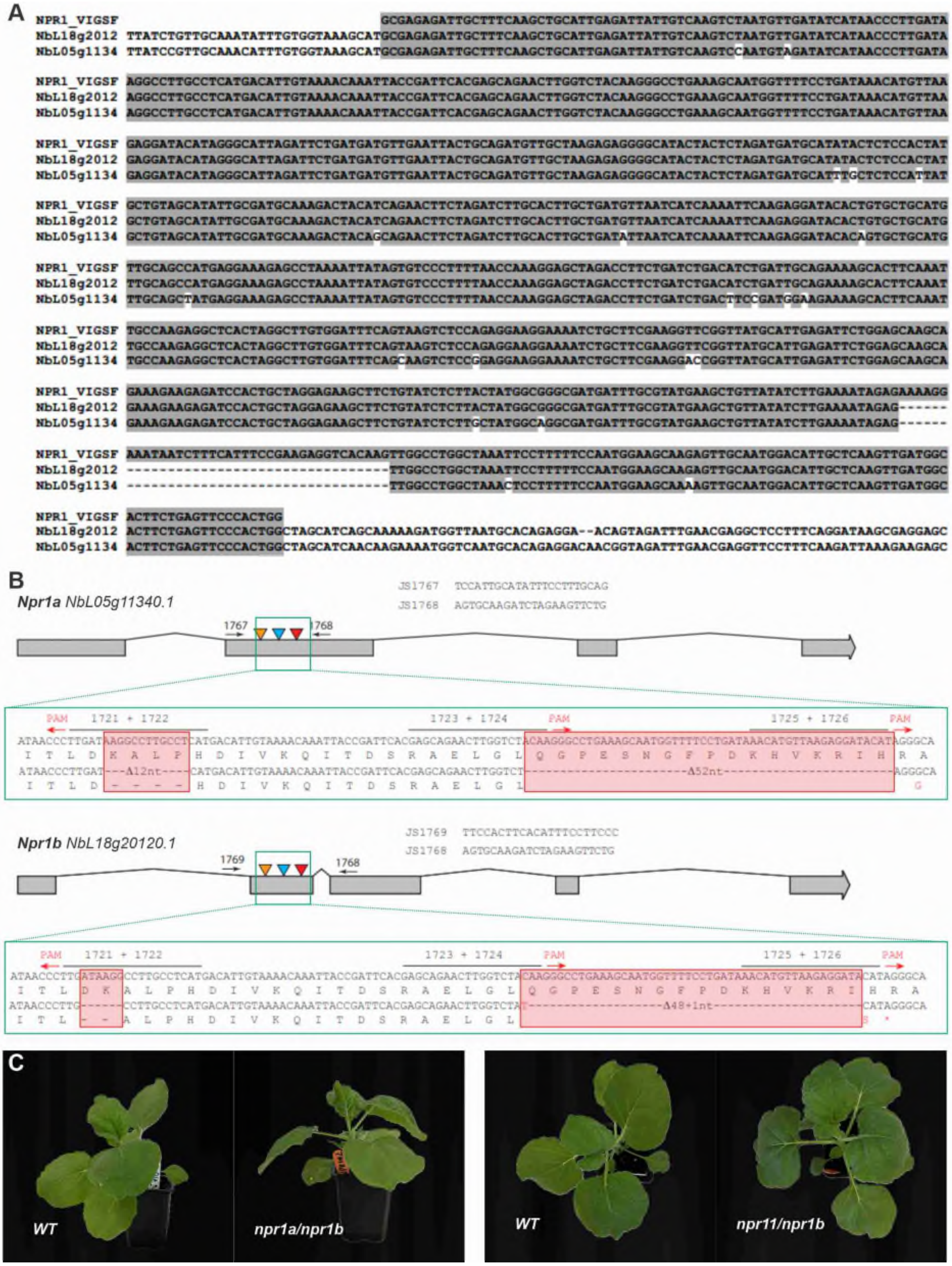

**Figure S6** The *npr1* double mutant of *N. benthamiana*.

**(A)** Alignment of the VIGS fragment (grey) targeting two different *Npr1* genes (LAB360 annotation). **(B)** Alignment of the genomic sequences of the two *Npr1* target genes of *npr1a/npr1b* mutant and wild-type *N. benthamiana*. **(C)** The *npr1#2* mutant has no growth phenotype. Pictures were taken of 4-week-old plants.

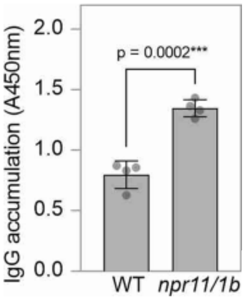


**Figure S7** Increased IgG accumulation in the *npr1a/npr1b* mutant plants.

The light and heavy chains of IgG COVA 2-15 were co-expressed with P19 and extracts were generated at 5 dpi and analysed with ELISA for antigen recognition. Error bars represent SD of n=4 replicates.

**Supplemental Table S1** Genes targeted by VIGS

| Gene | Function | Reference |
|---|---|---|
| ACIF1 | F-box protein acting downstream of immune receptors | Van den Burg et al. 2008 |
| AHA2 | Plasma membrane ATPase, alkalizes the apoplast | This work |
| BAK1 | Coreceptor kinase for receptor-like kinases (RLKs) and receptor-like proteins (RLPs) | Wang et al., 2018 |
| CERK1 | Receptor-like kinase for chitin | Wanke et al., 2020 |
| CDPK | Calcium-dependent protein kinase, activates RBOHB | Romeis et al., 2001 |
| CORE | Receptor-like kinase for recognition of cold shock proteins (CSPs) | Wang et al., 2016 |
| CRT3a | Calreticulin 3a, required for RLP/RLK/NLR folding | Liebrand et al., 2012 |
| EDS1 | Lipase-like protein, required for NLR function | Tran et al., 2016 |
| EIN2 | ER-localized transcriptional regulator activated by ET | Tran et al., 2016 |
| ICS | Isochorismate synthase; produces SA precursor | This work |
| MPK3/6 | MAP kinases -3 and -6, phosphorylate transcription factors | Duggan et al., 2021 |
| NPR1 | Transcriptional regulator of SA-responsive genes | Tran et al., 2016 |
| NRC2,3,4 | Helper NLRs for many immune receptors including RLPs and sensor NLRs | Wu et al., 2017 |
| PAL | Phenylalanine ammonia lyase; produces SA precursor | This work |
| RBOHB | Respiratory burst oxidase homolog, oxidative burst generator | Asai et al., 2008 |
| RE02 | Receptor-like protein recognizing secreted Cys-rich proteins, originally claimed to be the CSP receptor | Saur et al., 2016; Nie et al., 2021 |
| SAG101 | Lipase-like protein, interacts with EDS1 | Pombo et al., 2014 |
| SOBIR1 | Co-receptor kinase for receptor-like proteins (RLPs) | Liebrand et al., 2013 |
| WAK1 | Wall-associated receptor kinase | Rosli et al., 2013 |
| WRKY | WRKY-type transcription factors, controls transcription of immunity genes | Adachi et al., 2015 |

**Supplemental Table S2** Used plasmids

| **Plasmid** | **Description** | **Reference for plasmid** |
|---|---|---|
| pJK001c | Empty binary vector | Kourelis et al., 2020 |
| pJK-B2-022 | eGFP | Kourelis et al., 2023 |
| pID41 | *cGFP* | This work |
| pID42 | *sRFP* | This work |
| | *TRV::GUS* | Duggan et al., 2021 |
| | *TRV::PDS* | Liu et al., 2002 |
| pJK436 | *TRV::RBOHB* | This work |
| pID10 | *TRV::RE02 (/CSPR)* | Dodds et al., 2023 |
| pID24 | *TRV::CORE* | Dodds et al., 2023 |
| pID31 | *TRV::AHA2* | This work |
| pID32 | *TRV::CDPK* | This work |
| pID28 | *TRV::ACIF1* | This work |
| pID18 | *TRV::EIN2* | This work |
| pID15 | *TRV::NPR1* | This work |
| pID13 | *TRV::EDS1* | This work |
| pID21 | *TRV::WAK1* | This work |
| pID34 | *TRV::ICS* | This work |
| pID11 | *TRV::BAK1* | This work |
| pID14 | *TRV::SOBIR1* | This work |
| pID17 | *TRV::CRT3a* | This work |
| pID25 | *TRV::SAG101* | This work |
| | *TRV::NRC234* | Wu et al., 2017 |
| | *TRV::PAL* | This work |
| | *TRV::CERK1* | This work |
| | *TRV::MPK3/6* | Duggan et al., 2021 |
| | *TRV::WRKY* | This work |
| pDGE332 | | Stuttmann et al., 2021 |
| pDGE333 | | Stuttmann et al., 2021 |
| pDGE335 | | Stuttmann et al., 2021 |
| pDGE337 | | Stuttmann et al., 2021 |
| pDGE311 | 35S::zCas91 | Grutzner et al., 2021 |
| pDGE370 | Binary plasmid sgRNAs | This work |
| pJK268c | Binary P19 | Kourelis et al., 2020 |

| | | |
|---|---|---|
| pBM2 | Binary sfGFP | Mooney et al., 2025 |
| pKB21 | LC COVA2-15 IgG | Beritza et al., 2025 |
| pKB22 | HC COVA2-15 IgG | Beritza et al., 2025 |

**Supplemental Table S3** Used fragments for gene silencing

| **Length (bp)** | **Name** | **Sequence (5′ to 3′)** |
|---|---|---|
| 417 | *CDPK* | GAAGAAATTGCTGGTCTGAAAGAAATGTTCCGGATGATAGATACAGACAATAGTGGTCA AATAACCTTTGAGGAGCTCAAAGTTGGATTAAAAAGAGTCGGTGCTAATCTCAGGGAGT CTGAGATCTATGATCTTATGCAGGCAGCGGATGTTGATAACAGCGGCACAATTGATTAT GGGGAATTTATAGCAGCAACATTACATTTTAACAAAATTGAAAGAGAAGATCATCTATT TGCTGCTTTCTCTTACTTTGATAAAGATGGCAGTGGCTACATTACTGCAGACGAGCTTC AACAAGCTTGTGAGGAATTTGGCATTGGGGACGTCCATCTAGAAGATATGATAAGAGAC GCCGATCAGGACAATGATGGACGAATTGATTACAATGAGTTCGTTGCTATGATGCAAAA AGGA |
| 300 | *CORE* | TACAGACGAGTCACTGTTTTGGATCTTAGTTCTCTGAAACTGAGAGGTACTTTTTCACC AAGTATTGGAAATCTGAGCTTTCTTCGTGTCCTTGAGCTCCAAAATAATAGCTTTTCAG GTGAAATCCCATCAGAGCTTGGCTATTTGCATAAGTTACAGGTTTTACGCCTTGACAAT AACTCATTCACTGGTCATATTCCTTCAAACATTTCTGGTTGCTTCAACCTTTTTTCTGT TGTTCTTTCATATAATATGCTGGTAGGCATGATTCCAGCAGAATTAGGCACCTTGTCGA AACTT |
| 300 | *ACIF1* | TGTAAGAGTTTGAAGAAATTCTCATGTGGATCTTGCATGTTTGGAGCCAAAGGTATGAA TGCTTTGCTTGATCACTGTTCTACCCTTGAGGAATTATCTGTGAAGCGCCTACGTGGCA TCAATGATGGGTTTGCAGCTGACCCAATTGGCCCTGGAGCTGCGGCTTCATCGCTTAAA TCTATATGTTTAAAAGAGCTTTATAATGGCCAGTGCTTTGGGCCTTTGATTATTGGGTC CAAAAATTTGAGGACTTTGAAGTTGTTGCGATGTTTAGGTGATTGGGATAGGCTTTTTG AGACA |
| 300 | *AHA2* | GGCTGAAATTGCTAGGCAACGTGAGCTGCATACACTGAAAGGTCACGTCGAATCAGTGG TGAAGTTGAAAGGTCTTGACATAGAGACAATTCAGCAATCATACACTGTTTGAAAGGAG AGTATGTTCGACAAGTGTGATTTAACCGGTGAAGACATAATGAGGATGTGATTGTGAGT TACCTGTGGAAGAGACGAGAAATTAATAGTAGCAACATCTTAAAAAGTGCCTCATATAT TTGATATATATATATTCTTTAAGTTGGTCTTGCGTTTCCTATTTTCAAGTCTTGGCTCC ATGAG |
| 300 | *ICS* | AGTTTAAGATCTTGTCGTCCATTCACCCTACTCCAGCAGTTTGTGGGTATCCTACAGAA GATGCACGGGCTTTTATTTCAGAAACCGAAATGTTTGACCGAGGAATGTATGCTGGTCC TGTTGGTTGGTTTGGGGGGGAAGAGAGTGAATTTGCTGTTGGAATAAGGTCAGCTTTGG TTGAAAAGGGTCTTGGCGCATTAATTTATGCGGGAACTGGGATAGTGGAAGGAAGTGAT TCATCTCTAGAATGGGAAGAGCTCGAGCTGAAGACTTCACAGTTCACCAAATTGATGAA ACTTG |
| 383 | *GUS* | TAAAGAGCTGATAGCGCGTGACAAAAACCACCCAAGCGTGGTGATGTGGAGTATTGCCA ACGAACCGGATACCCGTCCGCAAGGTGCACGGGAATATTTCGCGCCACTGGCGGAAGCA ACGCGTAAACTCGACCCGACGCGTCCGATCACCTGCGTCAATGTAATGTTCTGCGACGC TCACACCGATACCATCAGCGATCTCTTTGATGTGCTGTGCCTGAACCGTTATTACGGAT |

| | | |
|---|---|---|
| | | GGTATGTCCAAAGCGGCGATTTGGAAACGGCAGAGAAGGTACTGGAAAAAGAACTTCTGGCCTGGCAGGAGAAACTGCATCAGCCGATTATCATCACCGAATACGGCGTGGATACGTTAGCCGGGCTGCACTCAATGTACACCGACA |
| 418 | *WAK1* | AGGCTACAAACAACTATGCCAGTGATAGAATTCTTGGTCGTGGTGGAAATGGAATTGTCTACAAAGGCATTCTATCTGATAATCGCATAGTTGCTATTAAGAAATCTAAGTTTATGGACGAGGAACAGGTTGAACAGTTCATTAACGAGGTACTTATTCTTACTCAAGTCAACCATAGAAATGTTGTGAGACTCTTCGGATGTTGTTTGGAAGCCGAAGTTCCTTTACTTGTCTATGAATACATTTCTCATGGAACTCTTTACGAGCATATCCACAATCGAAATGGAGCACCTTGGTTATCTTGGGAAAATCGGCTAAGAGTTGCTAGTGAGACAGCAAGTGCACTTGCTTACCTTCATTCATCCGCGCAAATGCCTATAATTCATAGAGATGTCAAGTCTGCCAATTTATTGTTGGA |
| 299 | *CSPR* | TGAAAAGTGAGAGATTTTTATTTCTCAATATTGCATTTTCAGCGTTCATCGGACTTGTTATTGGAACAAGTTCAGGAGGGGATGGTCGAACTACTTTGTGCATTGAGAGGGAGAGGGAAGCTCTTCTCAAGTTCAAGCAAGGTCTGATAGATAACTACGGTATCCTCTCGTCATGGGGGAGAGAAGAAGAGAAAGAAGAATGCTGTGGTTGGAAAGGTGTGCAGTGTAGCAATAGAACAGGTCATGTTGTGGTTCTTGATATTCATGCTCCGTCCTATAGTCAACATTTGAGAGGTAAC |
| 751 | *EIN2* | CTCTGATGGGAGAGCTCAATGGTATAATTCCATGGGATATGGACCGTCTGTTGGTCGATCAACGTACGAACAAGCCTATATGACTGGTTCACTGAGGGCAGGTGGTCCTCAGAGGTTTGAACATTCGCCATCTAAAGTCTGTAGAGATGCTTTCTCGTTGCAGTATAGCTCCTCCAATTCAGGGATTGGATCCGGATCCGGATCCCTGTGGTCTAGACAGCCTTTTGAGCAATTTGGTGTAGCTGGTAAGACGGATGTTGCTGCTAGCAGTGATCATGGAAATGTGCAGAGTTCATCTACTCCGGAGAGCACATCTACAGTCGACTTGGAAGCTAAGCTGCTTCAATCTTTCAGAAGTTGTATCGTCAAACTTTTGAAACTGGAAGGATCAGAGTGGTTATTTAGGCAAGATGATGGGACTGATGAGGATCTTATTGATCGGATTGCTGCAAGAGAGAAGTTTCTCTATGAGGCTGAAACTAGGGAGATAAGTAGATTGACCAACATTGGCGAATCGCATTTCTCTTCCAACAGGAAACCTGGTTCTGTTTCTGCTCCAAAACCTGAAGAGATGGATTACACCAAGTTCTTGGTGATGTCAGTTCCTCACTGTGGGGAAGGTTGTGTTTGGAAAGTAGATCTCATTGTTAGCTTCGGTGTATGGTGCATTCACAGAATTCTCGAGCTTTCACTTATGGAAAGTAGGCCAGAGCTGTGGGGCAAATATACCTATGTTCTCAACCGTCTTCAG |
| 788 | *NPR1* | GCGAGAGATTGCTTTCAAGCTGCATTGAGATTATTGTCAAGTCTAATGTTGATATCATAACCCTTGATAAGGCCTTGCCTCATGACATTGTAAAACAAATTACCGATTCACGAGCAGAACTTGGTCTACAAGGGCCTGAAAGCAATGGTTTTCCTGATAAACATGTTAAGAGGATACATAGGGCATTAGATTCTGATGATGTTGAATTACTGCAGATGTTGCTAAGAGAGGGGCATACTACTCTAGATGATGCATATACTCTCCACTATGCTGTAGCATATTGCGATGCAAAGACTACATCAGAACTTCTAGATCTTGCACTTGCTGATGTTAATCATCAAAATTCAAGAGGATACACTGTGCTGCATGTTGCAGCCATGAGGAAAGAGCCTAAAATTATAGTGTCCCTTTTAACCAAAGGAGCTAGACCTTCTGATCTGACATCTGATTGCAGAAAAGCACTTCAAATTGCCAAGAGGCTCACTAGGCTTGTGGATTTCAGTAAGTCTCCAGAGGAAGGAAAATCTGCTTCGAAGGTTCGGTTATGCATTGAGATTCTGGAGCAAGCAGAAAGAAGAGATCCACTGCTAGGAGAAGCTTCTGTATCTCTTACTATGGCGGGCGATGATTTGCGTATGAAGCTGTTATATCTTGAAAATAGAGAAAAGGAAATAATCTTTCATTTCCGAAGAGGTCACAAGTTGGCCTGGCTAAATTCCTTTTTCCAATGGAAGCAAGAGTTGCAATGGACATTGCTCAAGTTGATGGCACTTCTGAGTTCCCACTGG |

| | | |
|---|---|---|
| 323 | *BAK1* | AGCTCATAACTGGGCAACGGGCTTTTGATCTTGCTCGACTTGCAAATGATGATGATGTC ATGTTGCTAGATTGGGTCAAGGGACTTCTGAAGGACAAGAAGTATGAAACATTAGTAGA TGCAGATCTTCAAGGTAATTACGAAGAAGAAGAGGTGGAACAGCTTATTCGAGTAGCTC TTCTCTGTACAGGGAGCTCGCCGATGGAACGTCCAAAGATGTCAGAAGTGGTGAGAATG CTTGAAGGTGATGGCCTAGCTGAGAGGTGGGAAGAATGGCAGAAAGAGGAGATGGTCCG TCAGGATTATCCTGCTCACCACCCTCAC |
| 126 | *CRT3a* | ATGGCTCACTCTGAGCATAAACCAAGTAGACTGATATTCTCATTGGTACTAGTTTTGCT CTTCTTTACACTTGTCTCTTCTTCAGTATCTGAGATTTTCTTTGAAGAAAGCTTTGATG ATGGATGG |
| 327 | *SAG101* | TGCTAGGTCTTTTGGGGTTCTTCAACTTACTCCAAAGGCCAAGAATATGAGTCAGGATT CCTTGTTTAGTAGTGGCCAAGAATTGGCAAAGTTGGTGGTGAGCTCAGATCTACTGCAT GATTCTTGGGCTAGAAATTGTGATCTTCTTAATCATGCTTATTTGGATAATCCAACTAA CCCAGCTCCAATTGTGTTCAAAGTTTATTACCCATATTATACAAATGGTGCTATTGTTG CTTTTGTATCCTCCCCTACCTGTAGTATTCATCATCTTCAGAAAGAAATGGTCTCTTCA GAAGATCTTAAAGGTTCCCAAGTTGATTTTGA |
| 315 | *WRKY* | CTGTTGTGCTAATTCAGAAGAAGCATCACCTAAGAGGCCAAGGGAAATCAGAACCAATG TTGCAACTGTTTGTGTAAAAACTACTCCCTCCGATCAAAGTGCAATGGTGAAAGATGGA TATCACTGGAGAAAATATGGTCAAAAAGTCACAAGAGATAACCCTTCACCTAGAGCCTA CTATAAGTGTTCTTTTGCACCATCGTGCCCAGTCAAAAAGAAGGTGCAAAGAAGTGTAG AAGATCCATCAGTTTTAGCAGCTACATATGAGGGGGAGCACAACCATCCTCTCCCATCC CAAGCTCAAGTAACAGTGCC |
| 524 | *PDS* | AAGCTTGAAAATATCCACTGGAGCGGCAAACACAAAAGCATCTCCCTCGATTGCACTAC CGTCACTCAGTATAAAACTCTTGACACTTCCATCCTCATTCAGCTCAATCTTTTTTATT CGTGAGTTCAGTCTGACTTGGCCACCTTTTGACTCAATGTGTTCAACAATCGGCATGCA AAGTCTCTCAGGAGGATTACCATCTAAAAAGGCCATTTTTGAACCATGTTTCTCCTGAA GAAACCTGTCCAATGCGATCAAAATGCACTGCATTGAAAGTTCGTCAGGGTTTATAAAG TTGAGTGCCTTTGACATAGCAATGAACACCTCATCTGTCACCCTGTCCGGCACACCTTG CTTTCTCATCCAGTCCTTAACACTTATCCCATCTTGAGCTTCAACATAAGATTGCCCTC CAAGCATTGCTGGCAAGAGTCCAATTGCAAATTTGACTTTCTCTGGCCATGTAAGCATT TCGTTATTCTTTAAGATGGCTAAAATTCCATTTAAAGGAGCGGGTAAAGCTT |
| 193 | *RBOHB* | AATCATCATCCGCACCACCATCACCACCATTCGGACACAGAGATAATTGGAAATGACAG AGCGTCGTACAGTGGTCCGTTAAGCGGACCGTTGAATAAACGAGGCGGCAAAAAGAGTG CGAGATTTAACATTCCTGAATCTACCGACATCGGAACCAGTGTCGGAACCGGCGCCAAG TCCAATGATGATGCGT |
| 211 | *SOBIR1* | AATCTTTATCCACCAGATCATGCTGCACTTTTGCTTGTCCAAAAAGACTTGGGCATCCA AGGTCAACGCATTGCACTACTTTGCAACTCTGCAACAATATCCTGTGAAAGGCGAAAGG TAAACAGAACACAATTGTTGAGAGTCACCCGTATTGACTTCAGATCCAATGGATTGACT GGAACTTTATCTCCTGCCATTGGAAAACTTTCTG |
| 331 | *PAL* | AGCCGTGGACATCTTAAAGCTAATGTCATCCACATATCTAGTCGCACTTTGCCAAGCGA TAGACTTGAGGCATTTGGAAGAAAATCTGAGGAATGCAGTCAAGAACACGGTGAGCCAA GTCGCTAAGAGAACCTTGACGATGGGTGCCAATGGAGAACTTCATCCATCAAGATTCTG CGAAAAGGACTTGCTTCGAGTCGTGGACAGGGAATACGTCTTTGCATACGCTGATGACG CCTGCAGCGCTAACTACCCACTGATGCAGAAACTAAGGCAAGTCCTCGTCGACCACGCC TTAGAAAATGGCGAAAATGAGAAGGACGCAAACAGC |

| | | |
|---|---|---|
| 465 | *CERK1* | GTCTTGCTGCTGGCTGGTTTGGTTTACGTAGGATATTATAGGAAGAAAGCACAAAAGGT<br>TTCGCTGCTCTCTTCCGAAGACCGCCTCCATCAGTCTAGTCATGGCCCAGAGAGCAGCA<br>CGATAGTTAAAGCTGCAGATTCCGGTCGCCTGGCTAATGGCAATTCTCCAGAGCTTTCA<br>GGCATAACGGTGGATAAATCTGTTGAGTTCACTTATGAAGAGCTTGCTACTGCGACTAA<br>TGACTTCAGCATTGCGAACAAAATTGGACAAGGTGGTTTTGGTGCGGTTTACTATGCTG<br>AGCTCAGAGGCGAGAAAGCGGCTATCAAGAAAATGGACATGGAAGCTACAAGGGAGTTT<br>CTTGCTGAATTGAAGGTCTTGACACACGTTCATCACCTGAACCTGGTGCGCTTGATAGG<br>CTACTGTGTTGAAGGTTCCCTCTTCCTTGTATATGAATACGTTGAGAATGGC |
| 502 | *MPK3/6* | ATCACTACCAAGTATCGTCCTCCTATTATGCCTATTGCTCGTGGTGCTTATGGAATTGT<br>CTGCTCGGTGTTGAATACGGAGCTGAATGAGATGGTTGCAGTTAAGAAAATCGCAAATG<br>CGTTTGATAATTACATGGATGCTAAGAGGACTCTCCGTGAGATTAAGCTCCTCCGCCAT<br>TTAGACCATGAAAATGTAATTGGTTTAAGAGATGTGATTCCTCCACCGTTACGAAGGGA<br>GTTTTCTGATGTTTAGGACTTGAAGCCTAGCAATCTCCTCTTGAATGCCAACTGTGATT<br>TAAAGATATGTGATTTTGGGCTAGCTCGTGTCACTTCTGAAACTGACTTTATGACGGAA<br>TATGTTGTGACAAGATGGTATCGTCCACCTGAGCTATTGTTAAATTCATCTGACTATAC<br>TGCAGCAATTGACGTATGGTCAGTGAGTTGCATTTTCATGGAATTGATGGACAGGAAAC<br>CCCTATTTCCTGGTAGAGATCACGTACACC |
| 822 | *NRC234* | CACAAATCTGCCAACCCCTTTGTCCTTGTGAAGCTTAGCCTCAATCACAAACTTATCAA<br>TGGCATCTTCAGCAGAGTTGACCACAGTCTTAATTTTCTTCACCAATTCTTTGTGGACA<br>TCGTTCTCAGTGCGGGACTTAGCAGTTTGCTTGAGAAAAGCGTTGAAATCATTGAGATC<br>TTGAAGCAGACTCTCAGCCGAATCCTTAACTCCAACAATCAGCTCCGCATTGTCTCTTA<br>GCAGCTGCATCAAGTTCTGCACCAGAAACTCCACCGCAACGTTCGCCATACTGAGCAAA<br>TTTATTTTTATCGTCATGTCTTTTAGCTTCAATCACAAACTTATCAATTGAATCTTCAG<br>CATCATTTACCACTTTCCTTATCTTCTTTACTAGTGATTTCAAGACTTCATTGTCTCTC<br>CTTGATTTAGCTGCTTGTTTGAGAAAAGCATTGAAATCATTCAGATCTTGTAGTAAATT<br>TTCAACTTCACCCTTTATACCAATAATCAAGTCAGCGTTGTCGATTAATAATTGCATCA<br>AGTTTTCTACTAGGAATTTTACTGCTACATCTGCTGCTACATCTGCCATTTATTTTTCT<br>CTTTGTGCAGCTTCGCCTGAACCACAGACTTATCAACAGCATCCTCTGCGGTATGTACC<br>GTTTTTTGTATCTCCTCCACCAACACTTTCCACTGTTCGCTGTCGCTTGGTAATTTTGC<br>AGCATCGTCTAAGAACCCTTTTAGGTGTTGAACTTCTTTCAGCAGATTCTCTAACTCTC<br>CCTTTGCACTTCCGATCAGCTTTACGTTNTCAGTTAATAGCTGCAACAGGTTCTC |

**Table S4** Used oligonucleotides

| **Use** | ***name*** | **Sequence (5'-3')** |
|---|---|---|
| Cloning | *cGFPf* | TGGTGGTCTCAAATGCGTAAAGGCGAAGAG |
| Cloning | *cGFPr* | TTCGTGGTCTCAAAGCTCATTTGTACAGTTCATCCATACC |
| Cloning | *sRFPf* | TGGTGGTCTCAAATGAAGACTAATCTTTTTCTCTTTCTCATCTTTTCAC |
| Cloning | *sRFPr* | TTCGTGGTCTCAAAGCCTAGGCGCCGGTGGAG |
| Sequence | *TRV2f* | TGTTCAGGCGGTTCTTGTGTGTCA |
| Sequence | *TRV2r* | CCGTAGTTTAATGTCTTCGGGACATGC |
| RT-PCR | *CDPKf* | CTATGGTCCCGAAGCTGATG |
| RT-PCR | *CDPKr* | GGCCATGGATCTGATGAGAAGTC |
| RT-PCR | *ACIFf* | CCTTCACCCGAGCTCACTTC |
| RT-PCR | *ACIFr* | TTCATCGGGAAGCTCATATGTGT |
| RT-PCR | *COREf* | TGGCATTCGACAGTTTGGTG |
| RT-PCR | *COREr* | CAGACCCAAAACCACCCATG |
| RT-PCR | *ICSf* | ATTTCATGGTCCCTCAGGTTG |
| RT-PCR | *ICSr* | TCTGCACCCTCATAAGAACGG |
| RT-PCR | *CSPRf* | ACTGGGCTTTCCTCTTCTGC |
| RT-PCR | *CSPRr* | ACCCAAAGGCTTCAGGGATG |
| RT-PCR | *AHAf* | GTCTGAGTGCCGAGGAAGGA |
| RT-PCR | *AHAr* | CCCATGTAGTGTTCTCTGAGCAAG |
